# Temperate tree seedlings have similar drought vulnerability despite having different hydraulic drought responses in adults

**DOI:** 10.1101/2021.08.04.455116

**Authors:** Benjamin R. Lee, Inés Ibáñez

## Abstract

Climate change is projected result in higher frequencies of drought events across the world and lead to reduced performance in many temperate tree species. However, many studies in this area focus specifically on adult tree drought responses and overlook how trees in other age classes might differ in their vulnerability. Evidence shows that seedling drought response can differ from that of adults and furthermore that demographic performance in the seedling age class will have disproportionately strong effects on the assembly dynamics of future forests, together suggesting that understanding seedling drought responses will be critical to our ability to predict how forests will respond to climate change. In this study, we measured four indices of hydraulic response to drought (leaf water potential, photosynthetic capacity, non-structural carbohydrate concentration, and hydraulic conductivity), as well as interaction effects with shade treatments, for seedlings of two temperate tree species that differ in their adult drought response: isohydric *Acer saccharum* and anisohydric *Quercus rubra*. We found a strong isohydric response in *A. saccharum* seedlings that included conservation of leaf water potentials (>-1.8 MPa) and reductions in non-structural carbohydrate concentrations consistent with reduction of stomatal conductance. *Quercus rubra* seedlings were able to survive to more negative water potentials, but only rarely, and they showed a similar reduction in photosynthetic capacity as was found for *A. saccharum*. Our results suggest that, although *Q. rubra* seedlings display some anisohydric responses to drought, they are more isohydric than adults. Both species seem to be relatively similar in their vulnerability to drought despite the differences predicted from adult drought response, and our results suggest that seedlings of both species will be similarly vulnerable to future drought events.

## Introduction

Climate change is projected to increase temperatures and affect global patterns of precipitation, with many areas expected to become drier and hotter [1], and these environmental changes have the potential to strongly affect forest ecosystems [2]. Many studies have thus addressed the effects that water availability has on the performance of tree species, but primarily concentrating on adults [3–5]. However, in areas where projected climate may not significantly affect adult trees, relatively few studies address the effect that drought has on saplings and seedlings (but see Maguire and Kobe 2015, Kannenberg and Phillips 2020). This represents an important knowledge gap, as past research has demonstrated that drought response can significantly differ across ontogeny [8], suggesting that tree seedlings are likely to be affected by drought differently than their adult counterparts. Relatively small changes in water availability are likely to have profound effects on survival of younger life stages and could consequentially affect forest community assembly dynamics [2,9,10]. Scientists must therefore reconcile these differences to better predict the effects of climate change on forest demography.

Drought tolerance is a broad term that encompasses many plant traits including stomatal regulation behavior (e.g., iso/anisohydry), root morphology, xylem anatomy, and leaf abscission behavior [11–15]. Iso/ansiohydry, referring to whether plants close their stomata during drought to limit water loss [11], is an overarching concept that has been used to categorize a broad range of other drought-related traits. Plants are typically sorted along a gradient ranging from isohydric species which exhibit strong stomatal control on one end and anisohydric species which show little or no stomatal regulation on the other. This difference in behavior has physiological importance because prolonged stomatal closure (i.e., isohydry) can result in the over-depletion of labile carbon reserves caused by reductions in photosynthetic capacity, i.e., carbon starvation [16,17]. In contrast, anisohydric behavior can compromise the water column and result in catastrophic embolism and reduce xylem conductivity, i.e., hydraulic failure [11,18,19]. Anisohydric species are typically considered to be more tolerant of drought than isohydric species [11], but most species demonstrate some level of vulnerability to both carbon starvation and hydraulic failure due to their overlapping effects [12,20]. Moreover, a recent study found that plant vulnerability to drought is poorly predicted when using a singular plant hydraulic trait [21], and it has been further suggested that plant hydraulics should be studied and classified independently of the iso/anisohydry framework [22]. It is thus important that ecologists measure multiple drought responses to understand and then predict the consequences of drought.

There are several indicators of hydraulic strategy that can be used to partially explain a plant’s response to drought stress; these include regulation of internal water potentials [23], regulation of photosynthetic capacity [24], depletion of non-structural carbohydrates [16,25], and loss of xylem conductivity [26]. Predawn leaf water potential (Ψ_PD_) is a measure of plant hydraulic stress that is more representative of soil moisture conditions compared to midday leaf water potential, which is more strongly affected by hydraulic stress imposed by the atmosphere and is thus more negative [8,27]. Ψ_PD_ is especially sensitive to soil water availability in drought conditions for anisohydric species [28] and is therefore useful as a metric of plant water status in stressful conditions. Water potential has strong effects on the regulation of stomatal conductance [29], and stomatal regulation in turn affects Ψ_PD_ by allowing a plant to maintain internal water pressures above critical thresholds [11,18], below which the plant’s water column would snap, resulting in hydraulic failure. Tree species with strict stomatal regulation exhibit relatively narrow ranges of Ψ_PD_ that are maintained by reductions in stomatal conductivity until that threshold is surpassed and the plant dies [30]. In contrast, species that exhibit anisohydric behavior have much wider variation in Ψ_PD_ and can survive to much lower water deficits [30]. This strategy is common in species that have wide xylem conduits that can be refilled easily after water availability is reestablished [31] and species with physiological adaptations that prevent the spread of cavitation once it is initiated [32]. Strict regulation of Ψ_PD_ tends to be associated with species which are more vulnerable to drought [11], although there is recent evidence that this is not a fully generalizable rule [21].

Plant photosynthetic capacity (i.e., A_max_) is tightly linked to stomatal conductance, and thus plant water potential, during drought since stomatal closure limits CO_2_ from entering leaves [33]. This causes plants with strict stomatal regulation to reach net negative photosynthetic rates at less extreme soil water potentials compared to species that exhibit a more anisohydric drought response [11]. There is evidence of a strong relationship between stomatal regulation and changes in photosynthetic capacity across a wide range of species and biomes [23,24,34,35].

Without being able to rely on the assimilation of new photosynthate while stomata are closed, isohydric species instead mobilize labile carbon sources to meet plant energy demands [16]. This source of carbon, commonly referred to as non-structural carbohydrates (NSC), includes starch as well as soluble sugars such as glucose [25]. Reductions in NSC concentrations have been linked to drought [11,16,36] and other stressful conditions such as shade [6,37], and extreme reductions would theoretically result in death via carbon starvation, although it is unlikely that plants would need to exhaust their carbon supply to experience negative effects [16,38]. For example, refilling xylem following cavitation often requires the expenditure of energy [39] and soluble sugars are involved in plant osmoregulation such that reduced sugar content can lead to increased vulnerability to xylem cavitation [12,20]. Despite the difficulty associated with quantifiably measuring carbon starvation, the ability to increase or maintain [NSC] during drought is associated with drought-tolerant species that can afford to maintain stomatal conductivity due to other physiological and morphological traits [8,17].

Xylem conductivity (or hydraulic conductivity) is the rate at which fluid passes through a given stem or branch segment, and decreases with the amount of cavitation (i.e., gas embolisms) present within the segment [26,40]. Plants that conserve hydraulic conductivity through the closure of stomata are typically classified as isohydric and experience steep drop-offs in conductivity past a certain threshold [11]. Anisohydric plants are typically characterized by their more gradual decrease in conductivity that occurs across a wider gradient of water availability [11], a strategy that is commonly associated with drought tolerance. This strategy often coincides with wide xylem conduits (i.e., ring-porous anatomy), which are easier to refill following drought [31], and reduced stomatal regulation that provides the energy needed to expend in the refilling process [39].

Accurate classification of these traits is especially important for forest drought studies involving seedlings or saplings, which can substantially differ in hydraulic strategy compared to mature canopy trees. Cavender-Bares and Bazzaz (2000) showed that although adult *Quercus rubra* exhibit anisohydric behavior, seedlings were relatively isohydric, even when controlling for microenvironment differences between the two groups. The authors speculate that this is in part due to seedlings being unable to access deep water sources due to their shallow root profiles as well as to their higher vulnerability to environmental stress. Other recent research has also demonstrated that saplings received no tangible benefit from anisohydric behavior during drought and took longer to recover after the drought ended [41], suggesting that small size classes will have different vulnerability to drought even if they follow the hydraulic strategies of adult trees. Together, results from these studies support the recent call for a full quantification of hydraulic responses to drought [21,22] that is more robust in addressing suites of traits and within the context of ontogenetic differences.

Seedlings are the size class most likely to experience directional mortality effects [10,42,43], meaning that differences in species’ response to environmental drivers should be strongest in this phase of recruitment. Therefore, improving our understanding of ontogenetic differences in drought response will be critical to forecast changes to forest systems. Predicting future seedling performance will require an accurate estimation of hydraulic traits and behavior; if seedlings tend toward the same hydraulic strategies as their adult counterparts, climate change-related drought could accentuate differences in species performance based on these traits.However, if seedlings tend to be more isohydric than adult trees in general, as suggested by Cavender-Bares and Bazzaz (2000), drought effects could be more homogenous across species, increasing the relevance of other drivers of recruitment dynamics that affect community assemblage.

One such driver is light availability, which can severely limit photosynthetic rates even in species with high photosynthetic capacities [44]. Shade-tolerant species, i.e., species that can withstand prolonged periods in full shade, typically have relatively lower respiration demands than shade-intolerant species that allow them to persist on limited resources [45]. They may also exhibit phenological escape behavior that allows them to assimilate the resources they need in a short period of time when light availability is high [46–49]. Light limitations can lead to carbon deficits analogous to carbon starvation caused by stomatal limitations during drought [37,48] and can strongly shape recruitment in forest understories [50–52]. Light availability is also likely to affect seedling drought response [6,37]. Carbon starvation caused by deep shade can potentially compound with carbon starvation caused by stomatal limitations during drought [12,53]. However, there is also evidence that shading can ameliorate drought stress [54], particularly for drought-intolerant and shade-tolerant species [55]. These positive effects of shading are projected to drive seedling recruitment in some systems [56,57], and should therefore be considered alongside drought when studying the effects of climate change on forest demography.

In order to evaluate the extent to which drought response in tree seedlings differs from their adult counterparts, we measured four ecophysiological drought responses (leaf water potential, photosynthetic capacity, non-structural carbohydrate concentration [NSC], and xylem conductivity) for seedlings of two temperate tree species that commonly occur throughout eastern North America: isohydric *Acer saccharum* and anisohydric *Quercus rubra*. We used a greenhouse experiment to combine artificial drought and shading treatments. Our research questions were 1) Does seedling hydraulic strategy differ between species where adult hydraulic strategies are different? 2) How is seedling hydraulic strategy affected by light availability? And, 3) what are the implications for seedling demographic performance under climate change? We hypothesize that if seedlings of these species have hydraulic strategies similar to those used by their adult counterparts, then: 1) the range of leaf water potential experience by *A. saccharum* seedlings is much narrower than those observed in *Q. rubra* seedlings. 2) Under drought conditions, reductions in photosynthetic capacity are steeper in isohydric *A. saccharum* than in anisohydric *Q. rubra*. 3) The drop in NSC levels under drought is larger for *A. saccharum* seedlings. And, 4) the magnitude of the decrease in xylem conductivity when exposed to drought is higher for *A. saccharum*. Addressing these questions and hypotheses will contribute to our knowledge on how climate change will affect the demographic processes that shape temperate forest communities.

## Methods

We studied two temperate deciduous tree species that commonly co-occur across eastern North America, and that differ in their respective tolerances to shade and drought: *Acer saccharum* (Marsh.) and *Quercus rubra* (L.). *Acer saccharum* is a strongly shade-tolerant species [58] that is intolerant of drought [59]. It has relatively narrow xylem conduits (i.e., diffuse-porous xylem; Ellmore et al. 2006) and has been shown to exhibit isohydric stomatal behavior in response to drought [24]. In contrast, *Q. rubra* is moderately shade-tolerant [60] and moderately drought-tolerant [59]. Trees of this species have large xylem conduits (i.e., ring-porous xylem; Ellmore et al. 2006) and exhibit anisohydric stomatal behavior in response to drought [8]. Although wider xylem conduits are more vulnerable to embolism, there is also evidence that they are easier to refill following drought [31], conferring the higher drought tolerance generally associated with this trait. Adult *Q. rubra* trees also develop deeper roots than *Acer* species [61], which may enhance their relative drought-tolerance by allowing them to access deeper water sources.

Seeds of both species (from two different sources) were cold-stratified beginning in December 2016 and sown in large plastic tubs filled with potting soil (SunGro Horticulture; Agawam, MA, USA) at the Matthaei Botanical Gardens greenhouse (42.2996° N, 83.6630° W) the following spring. Once seedlings developed their first true leaves, we carefully removed them from the tubs and transplanted them into individual pots (volume = 313 cm^3^), supplementing new potting soil as needed. Sixteen seedlings of each species x source combination were randomly assigned to each treatment group (control, shade, drought, shade and drought) for a total of 256 total seedlings; 191 of these seedlings survived to be used in our analyses. Initial height for each seedling was measured two weeks following transplantation in order to account for maternal effects [57].

### Experimental Design

Following transplantation to individual pots, seedlings were immediately moved under moderate shade cloth (∼40% ambient PAR) and allowed to grow for an entire growing season (summer of 2017) under well-watered conditions. This was done to minimize the effects of first-year transplant stress and to allow seedlings the ability to assimilate enough carbon to allocate photosynthate to storage tissue and other labile carbon pools. Seedlings were moved to an outdoor pit following the onset of leaf color change in fall to allow seedling foliar phenology to respond to natural climate conditions. The pit was used to help insulate seedling roots from frost conditions they would otherwise not experience, and seedlings were further insulated by surrounding the pots with potting soil.

Seedlings were removed from the pit and moved back into the greenhouse in early spring 2018 corresponding to when leaf bud expansion was noted for both species. All pots were moved under one layer of shade cloth and were regularly watered until treatments were implemented. Environmental sensors were added simultaneously with when the seedlings were moved back inside, measuring air temperature and relative humidity (HOBO U23 Pro v2 data loggers) and light (HOBO Pendant data loggers; Onset Computer Corporation; Bourne, MA, USA). Temperature, relative humidity, and light were all measured at 30-minute intervals. Soil moisture was also measured at the individual (pot) level coinciding with harvesting or gas exchange measurement using a FieldScout TDR300 soil moisture meter (Spectrum Technologies; Aurora, IL, USA).

We took pre-treatment measurements of photosynthetic capacity and non-structural carbohydrate concentrations approximately four weeks following initial seedling leaf-out (June 14^th^), after which treatment conditions were initiated. Our four treatments were drought (D; no additional shade cloth added, seedlings are no longer watered), shade (S; extra shade cloth used to reduce PAR to 10% of ambient, seedlings remained well-watered), shade and drought (DS; extra shade cloth added, seedlings are no longer watered), and a control treatment (C; no additional shade cloth, seedlings remain well-watered). All seedlings acclimated to the study treatments for two weeks before three harvests were made at weekly intervals. Seedlings in each treatment combination were randomly assigned to be harvested for measurement of either xylem conductivity or [NSC]. Each harvest included six seedlings from each group: two for measurement of xylem conductivity and four for measurement of [NSC]. Predawn leaf water potential (Ψ_PD_) was measured on the morning of each harvest before sunrise as an approximation for soil water potential. Water potential was measured using excised leaves and a Scholander pressure chamber (PMS Instrument Company, Albany, OR, USA).

### Gas exchange measurements (A_max_)

The day before each harvest, for each treatment, we measured gas exchange in two of the four seedlings selected for the NSC analyses. We used an LI-6400 Portable Photosynthesis System equipped with a CO_2_ mixer assembly, LI-02B LED red/blue light source, and LI-06 PAR sensor (Li-COR Biosciences, Lincoln, NE, USA). We constructed light curves (i.e., *A-Q* curves) for each plant by recording gas exchange at 1500, 1000, 750, 500, 250, 100, 50, 25, and 0 μmol photons m^-2^ s^-1^ at CO_2_ concentrations of 400 ppm, ambient temperature, and ambient humidity. Maximum photosynthetic capacity (*A*_*max*_) was calculated using equations published by Marshall and Biscoe (1980) and using the *nls* command in the *stats* package in *R* v3.5.3. This parameter represents the maximum photosynthetic rate that a leaf is capable of under saturating light conditions. Reductions in *A*_*max*_ indicate limitations on the photosynthetic machinery, such as from reduced stomatal conductance [24].

### Non-structural glucose concentrations [NSC_Glu_]

After Ψ_PD_ measurement, tissue from seedlings selected for the NSC_Glu_ analysis were immediately separated into three pools: leaves (including petioles), stem (above root collar), and roots (below root collar). Leaves were microwaved for 180 seconds at 800 watts to stop leaf enzyme activity [25] and then all tissues were transferred to a drying oven and dried for 48 hours at 70 °C. We weighed each sample, ground them using a ball mill, and then stored the samples at 20 °C in airtight containers. We measured out 50 mg of each sample into screw-top conical tubes and soluble sugars were extracted according to Quentin et al. (2015) using repeated incubation and centrifuging in 80% ethanol. We measured glucose concentrations using a phenol-sulfuric acid colorimetric assay as described by Dubois et al. (1956). Glucose concentration was measured against a glucose standard curve using absorbance measured at 490 nm. Measurements were then converted to units of mg glucose per gram of dry tissue.

### Xylem conductivity (k)

Following Ψ_PD_ measurement, seedlings harvested for xylem conductivity measurements were removed from their pots and their root balls were soaked in water for 10 minutes to alleviate stress on the water column. We then cut stem segments from each plant underwater, recording the length of each segment and its average diameter. Conductance was measured using protocol established by Kolb et al. (1996). We slightly modified this protocol by using 20 mM KCl solution filtered to 0.22 μm (used to prevent microbial growth within the system that could cause artificial xylem blockages). Flow rate of the solution through the stem segment was measured at pressures of 0, -8, -16, -24, and -32 KPa. Hydraulic conductance (*k*) was calculated as the slope of the relationship between flow rate and pressure [26,64]. Flow rate was then standardized by segment length and cross-sectional area. Our vacuum source was not strong enough to forcefully remove embolisms from the stem segments and so we were unable to calculate the percent loss of conductivity.

## Analyses

Analyses were performed for each species independently. For each analysis we addressed our specific questions by trying different types of relationships between the variables involved (e.g., linear, exponential, additive, interactions), and tried several combinations of additional explanatory variables (i.e., initial seedling height and seed source). We describe below the models with the best fits based on Deviance Information Criterion for the photosynthetic capacity model (DIC; Spiegelhalter et al. 2002) and based on residual sum of squares comparisons for the other two analyses. Analyses used data from all harvests where seedlings from all relevant treatments were alive to be measured, with the exception being xylem conductivity measurements for *A. saccharum* where there were no data available for seedlings in the drought treatment.

### Predawn leaf water potential

Leaf water potential measurements (Ψ_PD_) taken over the span of the experiment (Fig 1) were used to assess the range of water potentials experience by seedlings of each species. Measurements were pooled across all seedlings harvested for both NSC and xylem conductivity experiments. We conducted a one-way ANOVA to assess whether Ψ_PD_ differed significantly between the two species.

**Fig 1.**
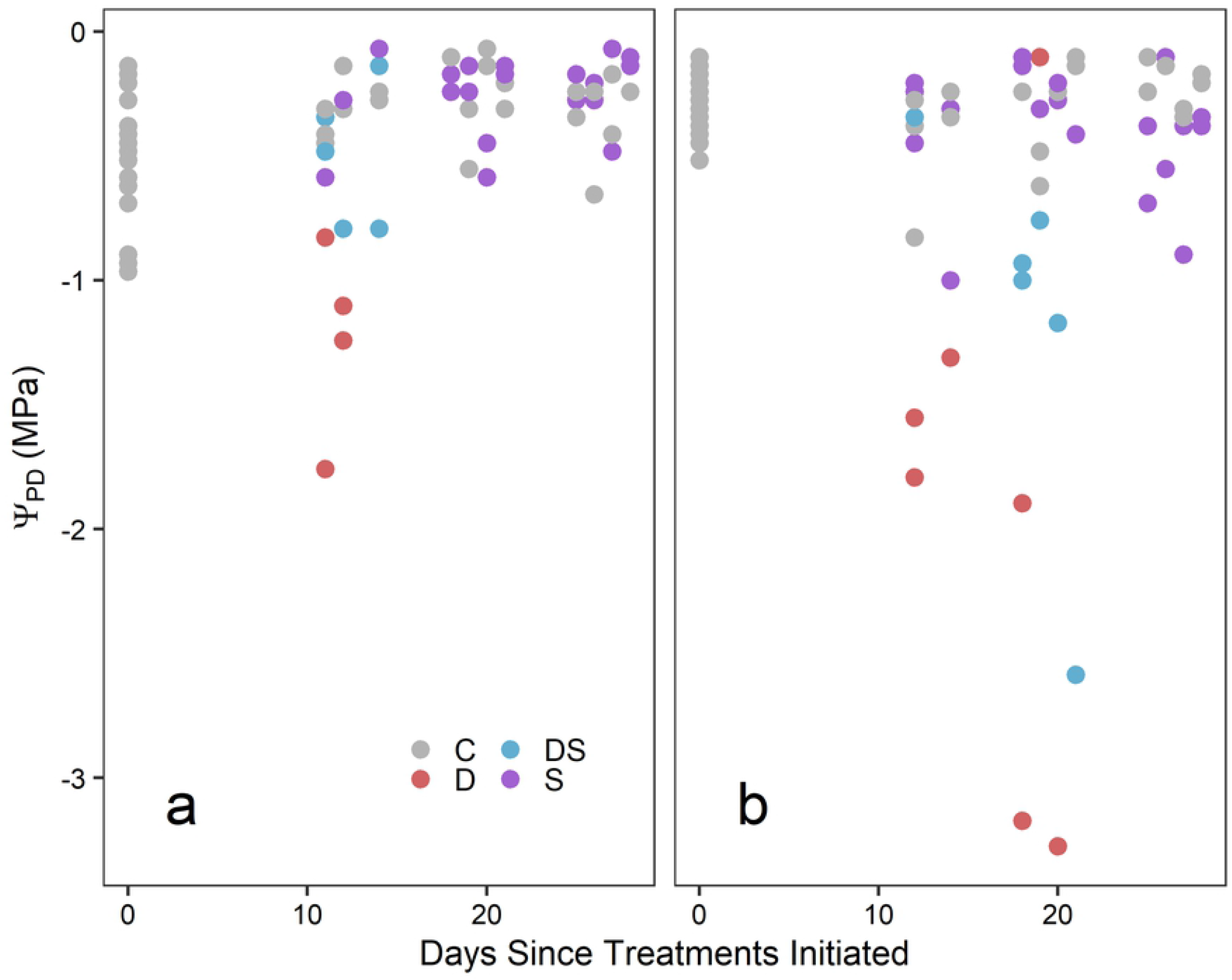
Predawn plant water potential over duration of the study. Predawn plant water potential (Ψ_PD_) plotted over the duration of the experiment for *A. saccharum* (a) and *Q. rubra* seedlings (b). Differences in color represent the different treatments: control (grey), shade (blue), drought (red), and combined drought and shade (purple).

### Photosynthetic capacity

We modeled maximum photosynthetic capacity (*A*_*max*_) as a function of light treatment and of leaf water potential (our proxy for drought effects); thus we combined seedlings from the two shaded treatments (S and DS) as well as from the two unshaded treatments (C and D) to assess the two light treatments and then used the full range of observed water potentials to account for drought treatments. Photosynthetic capacity for seedling *i* was estimated from a normal likelihood:

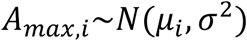

And an exponential process model that described well the reductions in A_max_ observed in the data (Fig 2a-b):

**Fig 2.**
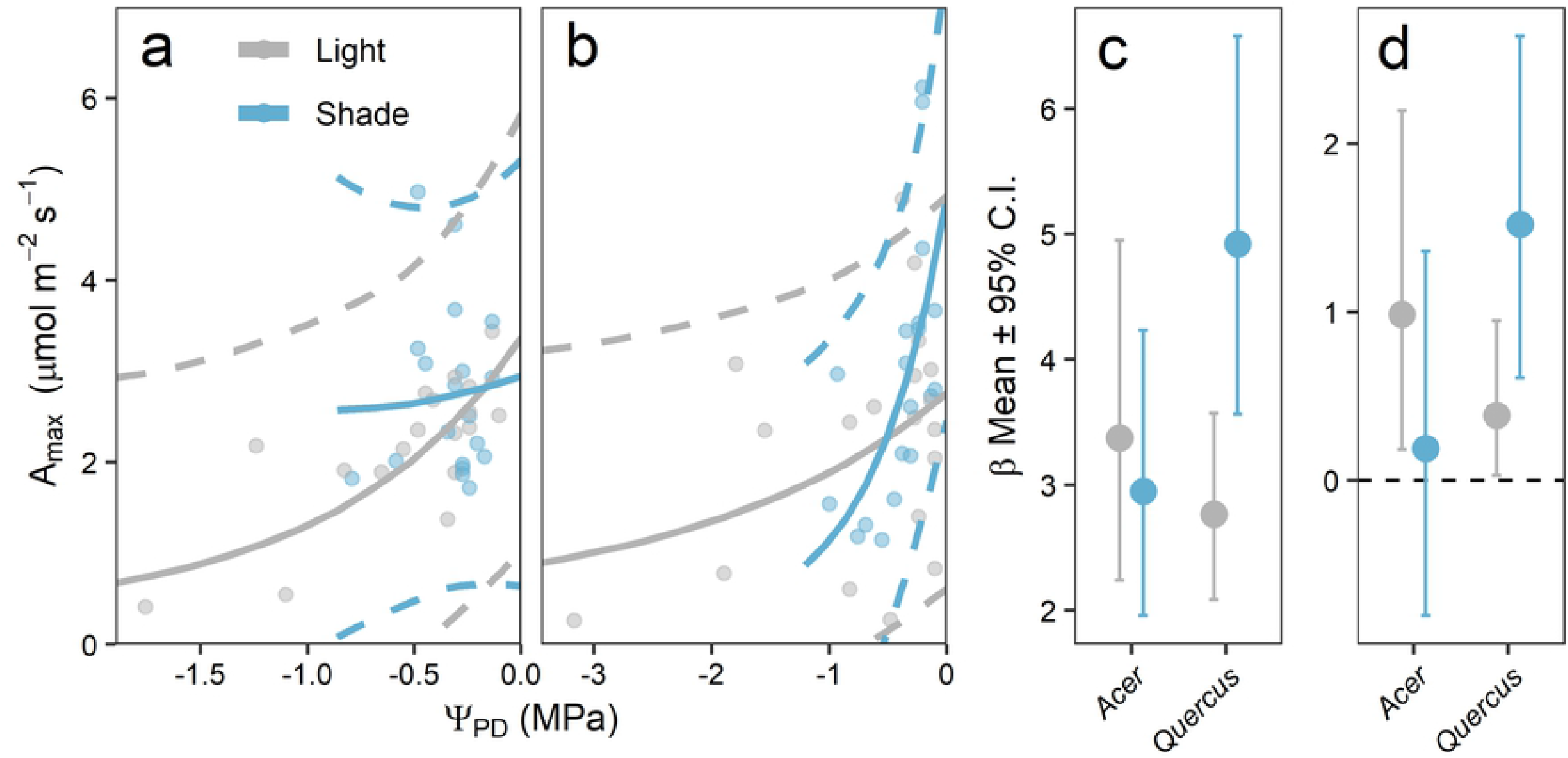
Relationship between Ψ_PD_ and maximum photosynthetic capacity. (a-b) Observed (points) and modeled (lines) relationship between Ψ_PD_ and maximum photosynthetic capacity (A_max_) for (a) *A. saccharum* and (b) *Q. rubra* seedlings. Colors represent seedlings in the non-shaded (grey) vs shaded (blue) treatments and dashed lines represent 95% predictive intervals. (c-d) Posterior estimated means (± 95% credible intervals) for (c) the intercept parameter β1 and (d) the decay parameter β2 relating Ψ_PD_ to A_max_. Parameters β2 are considered significantly positive if the 95% credible intervals (CI) do not overlap with zero and are considered different from each other if their 95% CI do not overlap.

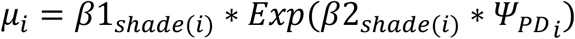

Parameter β1 represents the maximum photosynthetic rate at water potential equal to zero (i.e., full water availability). Parameter β2 indicates the decay rate at which *A*_*max*_ changes in response to changes in Ψ_PD_; we used this parameter to assess reductions in photosynthetic capacity. Both parameters were estimated for each shade treatment. The model was also evaluated for the effects of seed source and initial height, but they did not improve model fit and so we did not include them in the final model.

Parameter β1 was estimated from non-informative normal priors constrained to be positive, β1_*_ ∼ *N*(0,1000), and parameter β2 was estimated from non-informative uniform prior β2_*_ ∼ *Uniform*(−1,1). Model variance was estimated from a non-informative gamma prior distribution 1/ σ^2^ ∼*Gamma*(0.01, 0.01). We ran the model using two Monte Carlo chains for 40,000 iterations following a 60,000-iteration burn-in period. Analyses were completed using OpenBugs statistical software v3.2.3 [66] and model convergence was assessed using the Brooks-Gelman-Rubin statistic [67]. Parameter values (means, variances, and covariances) were estimated from their posterior distributions.

### Non-structural carbohydrates

Since seedling glucose concentrations did not change significantly over the duration of the harvest periods (Fig S1), data was pooled across harvest dates for which seedlings from all treatments were still alive (harvest 1 for *A. saccharum* and harvests 1 and 2 for *Q. rubra*). Non-structural glucose concentrations ([NSC_Glu_]) were analyzed using ANOVA that estimated the effects of treatment on values pooled by tissue type (leaf, stem, and root), or averaged across all pools (using averages weighted by the mass of each pool). Analyses were conducted separately for each pool x species combination. We only included data for seedlings that were recorded as alive at the time of harvest because most of the dead seedlings appeared to have died from hydraulic failure, and therefore could skew the results.

### Xylem conductivity

For each species, we analyzed the differences in xylem conductivity between seedlings harvested before and after the initiation of drought and shade treatments. We limited our analysis to compare differences only using seedlings harvested in the second harvest period (three weeks after the initiation of treatments) because there were no surviving seedlings in either of the two drought treatments past this harvest. We used initially carried out an ANCOVA to estimate the effects of treatment (C, D, S, DS), maternal effects (initial height), and of seed source. The effects of initial height and seed source did not improve the model fits and were therefore excluded from the final analysis.

Seedling conductivity and [NSC_Glu_] analyses were conducted in R (v3.5.3) using the *aov* and *anova* commands in the *stats* package to fit and compare models, respectively. Significant differences between treatments were estimated using Tukey’s HSD test, performed using the *TukeyHSD* command in *stats*.

## Results

Average daytime light levels across the two light treatments (Fig S2a) were consistent with light levels measured in related field experiments [48]. Light levels in the deep shade treatments were 30% (± 0.05% s.d.) of the light levels in the control light treatment. Average daily temperature (Fig S2b) was also consistent with field observations [48], although maximum hourly temperatures were much higher in the greenhouse than what has been observed in the field (Fig S3), and well above the temperature increases predicted under extreme climate change conditions for the Great Lakes region [68]. Temperatures were consistent across all treatments. We found some significant differences in height and mass between populations for each species (summarized in Appendix S1), but they did not significantly affect any of our analyses and are not included in the following results.

### *Predawn leaf water potential* (Ψ_PD_)

Leaf water potential decreased under the drought treatments for both species (Fig 1) with *Q. rubra* seedlings reaching more negative water potentials (−3.28 MPa) than *A. saccharum* seedlings (−1.76 MPa). However, we did not find any significant difference in Ψ_PD_ between the two species (Pr(>F) = 0.291). All *A. saccharum* seedlings in the D and DS treatments died before the second harvest (∼21 days after treatments were initiated) and all *Q, rubra* seedlings died before the third harvest (∼28 days after treatment initiation).

### Photosynthetic capacity

Our model fit (r^2^ of predicted vs observed) was 0.289 for *A. saccharum* and 0.305 for *Q. rubra*. Posterior parameter estimates can be found in Table S1. There were no significant differences in posterior estimates of intercept parameter β1 between light treatments for either species (Fig 2c), although values were higher in the shade treatment for *Q. rubra*. Decay parameters β2 were statistically significant, different from zero, for both shade treatments for *Q. rubra* but was only significant for the unshaded treatment in *A. saccharum* (Fig 2d). These decay parameters did not significantly vary between light treatments for either of the species. Predicted A_max_ decreased from 3.375 μmol m^-2^ s^-1^ (± standard deviation of 0.692) at Ψ_PD_ = 0 MPa to 1.309 ± 0.413 μmol m^-2^ s^-1^ at Ψ_PD_ = -1 MPa for unshaded *A. saccharum* seedlings, a decrease of 61.2% (Fig 2a). This drop was proportionally smaller in the shade treatment, with A_max_ decreasing from 2.936 ± 0.568 to 2.577 ± 0.929 μmol m^-2^ s^-1^ (12.3% decrease). This trend was the opposite for *Q. rubra* seedlings (Fig 2b), where predicted A_max_ decreased in light conditions from 2.775 ± 0.383 to 1.893 ± 0.325 μmol m^-2^ s^-1^ (31.8% decrease) and in shade conditions from 4.934 ± 0.746 to 1.137 ± 0.411 μmol m^-2^ s^-1^ (77% decrease).

### Non-structural carbohydrates

Non-structural glucose concentrations ([NSC_Glu_]) showed a general decrease over time across all treatments and pools for both species (Fig 3), but there were no significant differences between treatments for any of the species x pool combinations (Table S2). Some carbon pools significantly decreased from the pre-treatment values (Fig 3). There were no statistically significant drops in [NSC_Glu_] between control and treatment seedlings in either of the two species, and in some cases mean [NSC_Glu_] values were higher in the treatments than the control (particularly for *Q. rubra*, Fig 3e-h).

**Fig 3.**
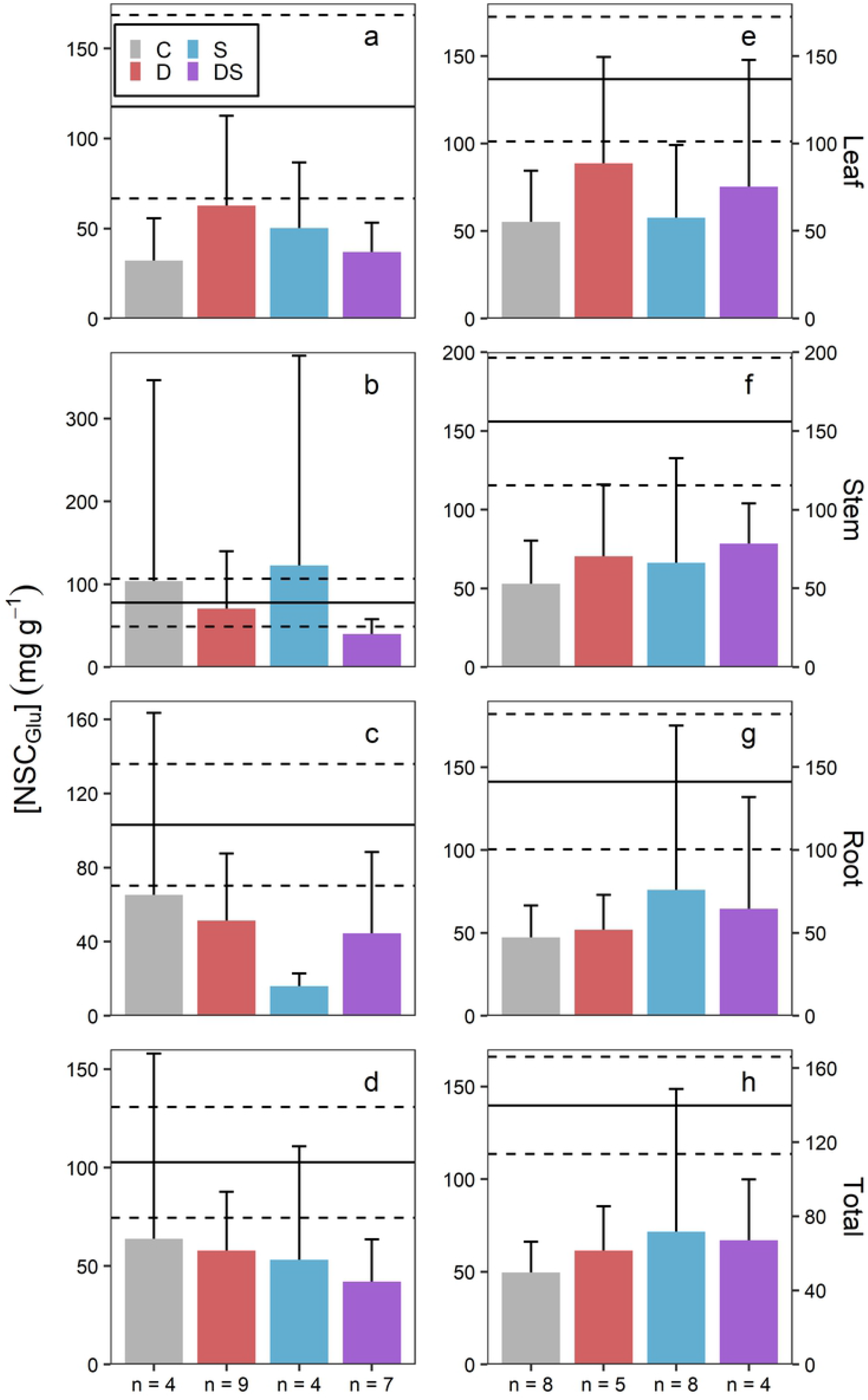
Non-structural glucose concentrations for seedlings in all treatment groups compared to pre-treatment controls. Non-structural glucose concentrations ([NSC_Glu_]; means + 2 s.d.) for *A. saccharum* (a-d) and *Q. rubra* seedlings (e-h) across the four experimental treatments. Rows show [NSC_Glu_] in leaf (a, e), stem (b, f), and root (c, g) pools as well as averaged across all pools and weighted by dry mass of each pool (d, h). Colors correspond to treatments as described in Fig 1. Horizontal lines indicate means (solid) ± 2 s.d. (dashed) of [NSC_Glu_] in seedlings harvested prior to the initiation of treatments.

### Xylem conductivity

There were no significant differences in conductivity between any of the treatments for either species (Table 1a; Fig S4). Average *A. saccharum* conductivity was 0.075 ± 0.016 g s^-1^ MPa^-1^ mm^-1^ (mean ± s.d.) in the control treatment and 0.073 ± 0.027 g s^-1^ MPa^-1^ mm^-1^ in the shade treatment (n = 4 in each treatment). There were no *A. saccharum* seedlings that survived in the drought treatments that could be used in this analysis. *Quercus rubra* average conductance was 0.076 ± 0.024 g s^-1^ MPa^-1^ mm^-1^ in the control treatment, 0.041 g s^-1^ MPa^-1^ mm^-1^ in the drought treatment, 0.082 ± 0.011 g s^-1^ MPa^-1^ mm^-1^ in the shade treatment, and 0.043 ± 0.021 g s^-1^ MPa^-1^ mm^-1^ in the combined drought and shade treatment (n = 3, 1, 4, and 3, respectively). We found a significant effect by drought treatment for *Q. rubra* seedlings when the different shade treatments were grouped together (Table 1b, Fig S5b), with seedlings of this species exposed to drought having significantly lower hydraulic conductivity (0.043 ± 0.017 g s ^-1^ MPa^-1^ mm^-1^) compared to those in the well-watered treatments (0.079 ± 0.021 g s^-1^ MPa^-1^ mm^-1^). *Acer saccharum* seedling conductivity in the combined well-watered treatments was 0.074 ± 0.021 g s^-1^ MPa^-1^ mm^-1^.

**Table 1:** ANOVA statistics describing (a) the differences in xylem conductivity between all four factorial treatments and (b) the differences between drought treatments for seedlings of both species. Drought treatment *A. saccharum* seedlings were not available to be used in this analysis and are thus excluded.

| (a) | Df | Sum Sq. | Mean Sq. | F value | Pr(>F) |
| --- | --- | --- | --- | --- | --- |
| *A. saccharum* | 1 | $7.5 \times 10^{-6}$ | $7.5 \times 10^{-6}$ | 0.015 | 0.906 |
| *Q. rubra* | 3 | $3.45 \times 10^{-3}$ | $1.15 \times 10^{-3}$ | 3.307 | 0.087 |
| (b) | Df | Sum Sq. | Mean Sq. | F value | Pr(>F) |
| *A. saccharum* | - | - | - | - | - |
| *Q. rubra* | 1 | $3.37 \times 10^{-3}$ | $3.37 \times 10^{-3}$ | 12.1 | 0.007 |

## Discussion/Conclusion

There is wide variation in the hydraulic strategies that plants use to avoid, tolerate, and recover from drought, and these strategies may be more nuanced than what is represented by broad categorization such as iso/anisohydry [22]. Previous research has demonstrated that common drought response indicators such as stomatal conductance regulation are often dependent on environmental context [22] and other trait axes [7,21], suggesting that adequate representation of drought tolerance requires the simultaneous quantification of multiple indicators. Furthermore, there is evidence that hydraulic strategy in tree species can vary substantially along ontogeny [8], and there is a sizeable knowledge gap in the scientific literature for how seedling drought response differs from that of adults. Whether seedling hydraulic strategies differ from adult hydraulic strategies will affect our predictions of forest demography under climate change [69].

In this experiment we measured the hydraulic strategies used by seedlings of two dominant tree species that commonly co-occur across a wide range of eastern North American forests, and that differ in their response to drought (as measured in adults). We quantified four commonly measured indicators of drought tolerance (leaf water potential, photosynthetic capacity, non-structural carbohydrate concentrations, and hydraulic conductance) over the duration of a greenhouse dry-down experiment. We used two different light treatments to investigate the potential interaction effects between shade and drought on tree seedling performance, since shade can both exacerbate carbon starvation via low light levels and/or ameliorate drought stress via decreasing temperature.

Our results indicate that, as predicted from adult characteristics, *A. saccharum* seedlings appear to be slightly more vulnerable to drought due to their inability to survive past relatively moderate reductions in soil water potential (Fig 1a), whereas *Q. rubra* were recorded to survive to considerably lower levels (Fig 1b). *Quercus rubra* seedlings were also able to survive longer in the drought treatments, but only by about a week. Still, we did not find statistically significant differences between species in any of the drought indicators, suggesting that seedlings of these two species are still similarly vulnerable to drought. However, due to the relatively small sample sizes used in this study, it is important to be cautious with these results and the conclusions drawn from them. Overall, our results suggest that the increased drought frequency predicted for the Great Lakes region [68] could negatively affect recruitment and demography of both species in approximately equal measure.

### 1) Does seedling hydraulic strategy differ between species where adult hydraulic strategies are different?

Tree performance during drought is strongly affected by stomatal regulation via two interacting processes [11,20]. Restricting stomatal conductance allows trees to avoid excessively low internal water pressures that can embolize xylem and lead to hydraulic failure [18]. However, reducing conductance comes at the cost of reducing photosynthetic capacity, which makes it necessary for trees to consume labile carbon pools and increases the risk of dying from carbon starvation [16]. Trees exhibit variation in other traits such as rooting depth [70,71] and xylem pit anatomy [32] that can help mitigate the negative effects associated with one process or the other (e.g., by giving trees access to more resources or by helping them resist cavitation), but there is strong evidence that tree performance in drought conditions is strongly determined by tradeoffs between these two processes [20]. Furthermore, juvenile trees may not be able to make use of the same mitigating strategies used by adults due their size and relative lack of access to resources (i.e., deep water sources; Cavender-Bares and Bazzaz 2000). Hydraulic outcomes at the seedling level may therefore be more similar among species than they are at larger size classes.

Our results do not fully support the idea that seedling hydraulic strategies are similar to those of the adults; while leaf water potentials reached lower levels in *Q. rubra* seedlings, as we expected, the other responses to drought did not differ between the two species. First, while adult *Q. rubra* (as well as other *Quercus* species in general) respond to drought stress by maintaining photosynthetic capacity at the cost of reduced leaf water potential [8,24], we found that seedlings of this species exhibited declines in photosynthetic capacity that began at Ψ_PD_ < -1 MPa (Fig 2b). This was not significantly different from the trend found in *A. saccharum* seedlings (Fig 2a), which matched the photosynthetic response demonstrated in adults of this species [24]. This relatively isohydric response of *Q. rubra* seedlings agrees with previous work done by Cavender-Bares and Bazzaz (2000) and provides support for the idea that seedling hydraulic strategies will be more similar between species than they are in conspecific adults.

Adults of these two species have also been shown to have different wood densities, with *A. saccharum* having dense wood and diffuse-porous xylem and *Q. rubra* having ring-porous xylem and wood that is less dense [14]. Diffuse-porous xylem can help trees avoid embolism formation due to the narrower conduits, but they are also more difficult to refill after embolism occurs [31]. Narrow xylem conduits and strong stomatal control help *Acer* species maintain hydraulic conductance during drought [72,73]. In contrast, *Quercus* species are more prone to gradual but significant declines in conductivity [40,73,74]. We hypothesized that *A. saccharum* seedlings would demonstrate stricter control of internal water potentials compared to *Q. rubra*, and thus show little variation in the conductivity of living seedlings, whereas *Q. rubra* seedlings would show a gradual decline in conductivity associated with anisohydric stomatal regulation.

We found tentative support for this hypothesis with respect to *Q. rubra* seedlings, for which conductivity appeared to decrease gradually beginning at Ψ_PD_ < -1 (Fig 4), which closely resembles the pattern seen in conspecific adults [74] and agrees with previous work done on seedlings of this species [40]. However, we were not able to fully quantify this trend in a more complex analysis because we did not measure percent loss in conductivity (*sensu* Kolb et al. 1996) and because we had limited survival in our seedlings. There was high mortality for this species past the -1 MPa threshold, suggesting that these seedlings are still vulnerable to drought. *Acer saccharum* seedlings supported the hypothesis, with conductivity maintained at Ψ_PD_ > -1 and hydraulic failure past that point (Fig 4), which agrees with the strategy used by adults. We observed seedlings of this species surviving to slightly more negative Ψ_PD_ in the NSC harvests (Ψ_PD_ =-1.76 MPa, Fig 1), suggesting that reductions in conductivity may follow a trend more similar to that of the *Q. rubra* seedlings, but conductivity was not quantified for these individuals and thus we lack the evidence needed to better support this conclusion.

**Fig 4.**
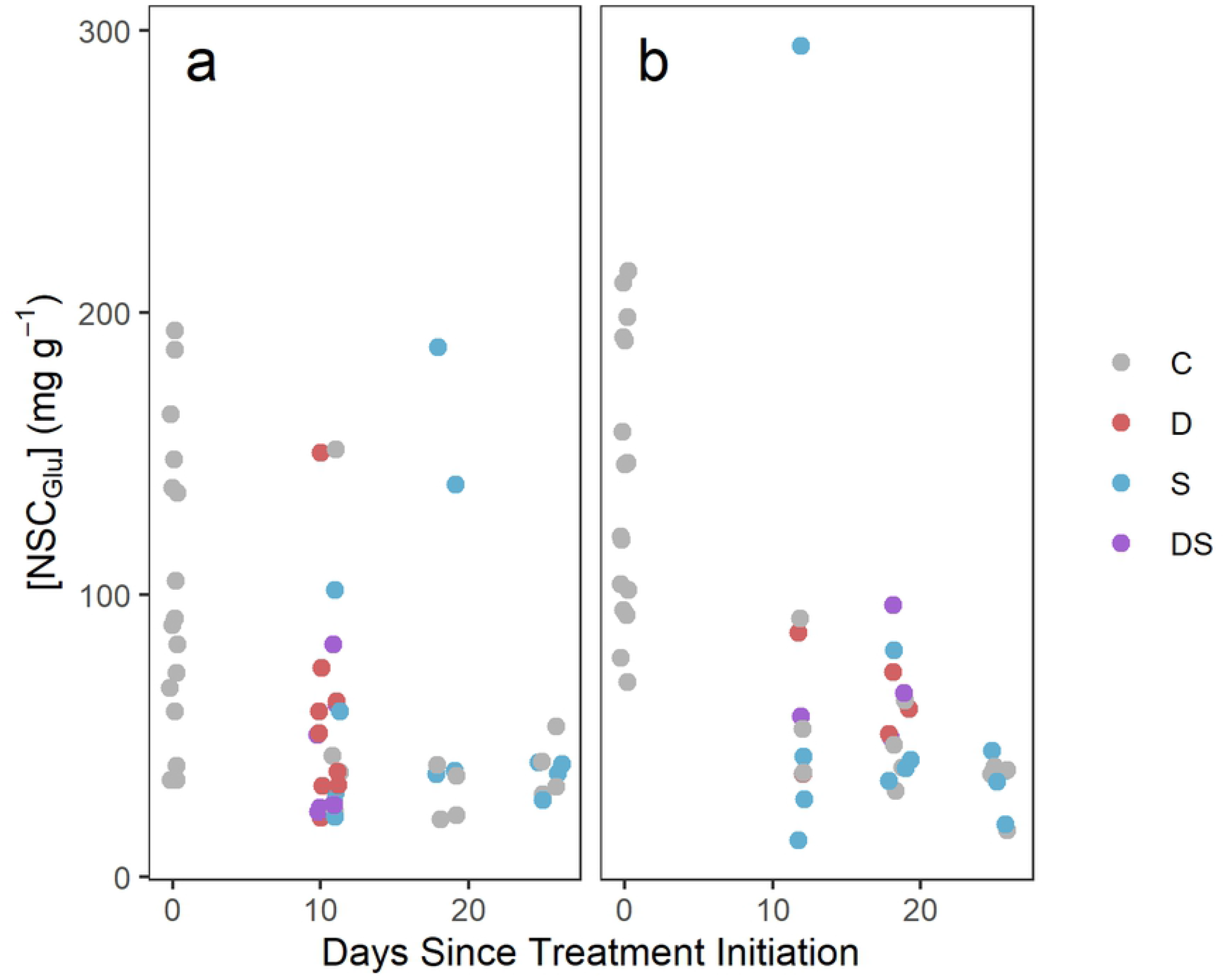
Xylem conductivity for both species in relation to predawn plant water potential. Xylem conductivity (k) for (a) *A. saccharum* and (b) *Q. rubra* seedlings plotted by predawn plant water potential (Ψ_PD_). These plots show data from across all three harvests for seedlings that were recorded as alive at time of harvest. Colors correspond to treatments as described in Fig 1.

Photosynthetic capacity and hydraulic conductivity are both strongly intertwined with changes to non-structural carbohydrate concentrations. Reductions in photosynthetic capacity limit the production of carbohydrates and make plants more dependent on labile carbon pools, reducing [NSC] to maintain metabolic rates [16]. Conductivity directly affects a plant’s ability to transport carbohydrates from source tissue (leaves) to tissues where sugars are needed (stems and roots). However, [NSC] also affects conductivity since energy is often required to refill xylem following cavitation [39] and because soluble sugars are necessary for osmoregulation processes [12,20]; thus, reduced sugar content can lead to faster hydraulic failure.

We found no significant differences in non-structural glucose between treatment conditions for *A. saccharum* in any of the tissue pools, and limited differences between treatment and pre-treatment [NSC_Glu_] (Fig 3a-d). This suggests that seedlings were generally able to maintain their labile glucose pools throughout the experiment. This is consistent with a previous study that worked with *Acer rubrum* seedlings (which are closely related to *A. saccharum* and typically exhibit a similar hydraulic strategy; Davies and Kozlowski 1977), which found reductions in soluble sugar concentrations over time in similar treatments [6]. We also found no significant differences between treatments in any of the *Q. rubra* NSC_Glu_ pools (Fig 3e-h), but there were more instances of [NSC_Glu_] reductions relative to pre-treatment controls. This contradicts previous research by Maguire and Kobe (2015), which found general increases in soluble sugar concentrations for this species under similar drought and shade treatments and suggests that seedlings of this species experienced similar stress across all treatments.

The lack of a significant difference between our control treatments and the three stress treatments prevents us from being able to make strong conclusions about how drought will affect [NSC_Glu_] for seedlings of either species. The difference between our results and the results from other studies that have found significant effects in similar experiments (e.g., Maguire and Kobe 2015) could be due to the extremely high temperatures experienced by seedlings in the greenhouse environment (Fig S3), which affected all seedlings equally and could have caused reductions in [NSC_Glu_] associated with high respiration rates [76]. It is also possible that the trends we report do not tell the full story as we did not measure starch concentrations, nor did we measure the concentrations of other soluble sugars besides glucose. Recent evidence suggests that soluble sugar concentrations can be maintained through the conversion (and therefore reduction) of starch [7], suggesting that measurement of total non-structural carbohydrate concentrations may tell a more complete story.

### 2) How is seedling hydraulic strategy affected by light availability?

There is experimental evidence for both the mitigating [37,54] and exacerbating effects [6,12,53] that shade can have on tree performance during drought. In temperate North American forests, understory light availability under full canopies can be 2-3 orders of magnitude lower than light availability in open canopy conditions [48], and access to light plays a significant role in tree seedling demographic performance in these systems [77,78]. It is therefore important that projections of tree recruitment under climate change account for any interactions between drought and shade.

Contrary to other tree seedling studies (e.g., Piper and Fajardo 2016), we did not find any significant effects associated with light availability in any of our drought response indicators. Although including light treatments improved our photosynthetic capacity model’s performance, there were no significant differences between light condition effects for either β1 (photosynthetic capacity at saturating water availability) or β2 (the decay rate parameter) (Fig 2c-d). There were no significant shade effects in any of the glucose pools (Fig 3) and hydraulic conductance was better predicted in an analysis that explicitly omitted light treatment and focused solely on differences in drought treatment (Fig S5). The lack of a shade effect on glucose concentrations is consistent with previous research that found no significant change in soluble sugar concentrations over time in seedlings [6] and saplings [7] of temperate tree species common in our study region. However, both studies found significant reductions in starch concentrations that suggest that plants prioritize the mobilization of starch for use in metabolism over the consumption of soluble sugars, potentially due to the importance of soluble sugars in osmoregulation [12,20]. We did not measure starch concentrations, so we can only speculate that this mechanism may have affected our results. The extreme temperatures experienced in the greenhouse could have also created respiration demands across all treatments that overwhelmed any signal that we might have otherwise observed with differences in light availability [76]. Still, our results do not support the existence of either mitigating or exacerbating effects of shade on hydraulic performance of temperate tree seedlings.

### 3) What are the implications for seedling demographic performance under climate change?

Altogether, the results from our experiment suggest a convergence in the vulnerability of temperate tree seedlings to drought that is inconsistent with drought responses expected from evidence collected in adult trees of the same species. There were no drought treatment *A. saccharum* seedlings that survived longer than two weeks past the initiation of experimental treatments and no water-limited *Q. rubra* seedlings that survived past three weeks. Although our stress treatments were harsh (and also likely exacerbated by high temperatures) the lack of a significant difference between seedling responses is consistent with results from previous research [6]. Still, the relatively small sample size of seedlings surviving in this experiment could have affected our ability to pick up on differences between the two species at finer scales of time, temperature, and soil moisture, and it therefore prevents us from making stronger conclusions about seedling drought dynamics.

Even so, with no clear distinction in seedling vulnerability to drought, potential differences in seedling demography and recruitment between species within the context of climate change are more likely to be driven by other factors. For example, we have previously shown that access to spring light [48] and the capacity to track climate change with spring leaf out phenology [47] differs between species and that this mechanism can help account for carbon starvation dynamics expected under hotter and drier summers. The lack of a difference between species further indicates that seedling drought vulnerability is decoupled from adult vulnerability and suggests that future community assembly in temperate forests will not be strongly limited by the vulnerability of tree seedlings to drought. This places a stronger emphasis on traits and behaviors that do differ between species and that could lead to differential outcomes during climate change.

## Acknowledgements

I. Ibáñez was funded by the NSF (DEB 1252664) and B. Lee was funded by the (Shrank Summer Research Support Fund). We thank D. Zak for the use of an IRGA for gas exchange measurements; DZ, R. Upchurch, and K. Wood for guidance and assistance on NSC analysis; DZ and D. Goldberg for providing valuable feedback on preliminary drafts; and D. Peltier for advice and guidance on modeling gas exchange measurements.

## Supporting Information Captions

**S1 Appendix 1**. Supplemental information concerning initial seedling height.

**S2 Supplemental tables and figures**. Tables S1 and S2 and Figures S1-S7

